# The Dominant Hand-Area Network: Task-Signal Mismatch and the Decoupling of BCI Skill from Global Efficiency in Postural Motor Imagery

**DOI:** 10.1101/2025.11.23.690036

**Authors:** Md Ahasanul Al Hasib Ayon

**Affiliations:** Dhaka Medical college

**Keywords:** Motor Imagery, Brain-Computer Interface (BCI), Network Neuroscience, Electroencephalography (EEG), Graph Theory, Granger Causality, Prefrontal Cortex, Sensorimotor Rhythms, Global Efficiency, Network Neuroscience

## Abstract

**Rationale:** The neurophysiological basis of motor imagery (MI) is foundational to Brain-Computer Interface (BCI) development. While BCI decoding relies on local sensorimotor rhythms (SMRs), MI itself is a large-scale network process. It is an open, and clinically critical question whether an individual’s BCI decoding skill is related to global network integration. This is complicated when the instructed task (e.g., postural imagery) differs from the underlying neurophysiological signal (e.g., hand-area SMRs). This study investigates the network dynamics of this task-signal mismatch and its relation to BCI performance.

**Methods:** We analyzed a 64-channel EEG dataset of 32 healthy participants (from the OpenNeuro dataset ds005342 cohort) performing a cued motor imagery task (e.g., “sit-to-stand”) contrasted against an idle state. BCI decoding performance was quantified using a Common Spatial Patterns (CSP) pipeline with Linear Discriminant Analysis (LDA), Support Vector Machine (SVM), and Random Forest (RF) classifiers. We then constructed functional brain networks from the beta (13-30 Hz) band using Phase-Locking Value (PLV) and spectral Granger Causality (GC). Graph theory was employed to analyze network topology, and we correlated individual BCI accuracy with global network efficiency in both sensor and source space.

**Results:** BCI performance was robustly high (LDA (Mean: 82.10% ± 8.80% SD)), 95% CI [78.92%, 85.28%], p < 10^−20^), confirming a high-fidelity local SMR signal. Strikingly, CSP analysis revealed this signal originated not from the expected leg-area (Cz), but from the hand-area sensorimotor cortex (C3/C4) (See **Figure 4**). Despite this local-signal fidelity, this decoding skill was critically decoupled from whole-brain network integration, showing *no* correlation of BCI accuracy with global efficiency, in either sensor space (r (30) = 0.189, p = 0.299) or, more critically, in source space (r (30) = −0.054, p = 0.769, 95% CI [−0.395, 0.300]). Our network analysis, centered on this C3/C4-dominant signal, revealed a novel, hypothetical M1-centric model: the motor cortex (C3/C4) acted as the primary information broadcaster (via GC), driving activity in prefrontal coordinating hubs (F3/Fz). Furthermore, the MI state was characterized by a dynamic reconfiguration, increasing the information-gating centrality of the premotor (FC1) node while suppressing the parietal (P3) node.

**Conclusion:** BCI decoding skill reflects a local process driven by SMR signal fidelity from the dominant hand area—even during non-hand tasks—rather than a global process relying on whole-brain efficiency. We propose a hypothetical “Broadcasting Motor Cortex” model in which the MI network is not a simple PFC–M1 hierarchy but a dynamic, M1-centered system that re-routes information through premotor and parietal gateways. Within this framework, baseline network properties may influence skill, but task-evoked global integration does not emerge as a strong correlate of performance. Given that our study was powered only to detect large correlations (r > 0.48), the absence of such effects suggests that baseline network states (16, 17), rather than active global integration, are more likely to predict individual BCI ability.

## 1. Introduction

### 1.1 Motor Imagery as a foundational Cognitive and Clinical Tool

Motor Imagery (MI) is the cognitive process of mentally simulating a motor action without any overt muscle contraction (1, 2). This “action simulation” engages a vast cortical network that largely overlaps with the one recruited during actual motor execution, including the primary motor cortex (M1), premotor cortex (PMC), and supplementary motor area (SMA) (3). This fundamental property has established MI as a powerful paradigm for understanding the neurophysiological basis of motor planning, volitional control, and neural plasticity. This understanding has formed the basis of modern translational neuroscience, particularly in the development of Brain-Computer Interfaces (BCIs) for neuro-rehabilitation (4). By decoding the *intent* to move directly from cortical signals (5), MI-BCIs provide a non-muscular channel for communication and control, offering a viable pathway for restoring function to patients with severe motor deficits (5, 6).

### 1.2 The Canonical Model: SMRs and Localized BCI Decoding

The “canonical model” of MI-BCI is built upon a well-established, local neurophysiological phenomenon: the modulation of sensorimotor rhythms (SMRs) (7). SMRs are endogenous oscillations prominent over the sensorimotor cortex, typically observed in the mu (8-12 Hz) and beta (13-30 Hz) frequency bands. The act of imagining movement induces a focal, contralateral decrease in the power of these rhythms, a phenomenon known as Event-Related Desynchronization (ERD) (1, 8). For decades, BCI research has leveraged the fact that imagery of *different limbs* (e.g., left hand vs. right hand) produces topographically distinct ERD patterns, most notably over the C3 and C4 electrode sites (9). This reliable, high-variance local signal is the primary, and often exclusive, feature used by the most successful BCI decoding algorithms (10). The Common Spatial Pattern (CSP) algorithm, in particular, is a spatial filtering technique designed specifically to find optimal projections of multi-channel EEG data that *maximize* the variance of this local signal for one class while minimizing it for another (11). Consequently, the success of an MI-BCI is almost entirely defined by the signal-to-noise ratio and fidelity of these local SMRs.

### 1.3 The Network-Level “Gap”: Is BCI a Whole-Brain Phenomenon?

While the BCI field has successfully treated MI as a *local signal-processing* problem, the field of cognitive neuroscience views it as a complex, *whole-brain network* process (3). This distributed network is widely assumed to be a top-down executive function (12, 13). In this conventional hierarchical model, the prefrontal cortex (PFC) provides the executive “command,” attentional resources, and working memory required to initiate and sustain the imaginary state (12). This command is thought to propagate to the PMC and SMA for motor planning, which in turn *drives* M1 to generate the final, simulated motor plan (13).

This discrepancy creates a critical, unbridged gap in the literature. The BCI field treats M1 as the *origin* of the decodable signal, while the cognitive field often treats it as the *recipient* of a prefrontal command. This leads to a pervasive, “common sense” hypothesis: that these two scales of analysis are linked. It is an implicit assumption that a “better” or more efficiently organized global brain network—one with high global efficiency (14, 15)—should lead to “better” cognitive (and thus BCI) performance. It remains a fundamental, and surprisingly untested, question whether an individual with a more highly integrated and efficient global brain network (17) is also better at generating the specific local signal required for BCI control.

### 1.4 Our Solution: A Dual-Analysis Framework

This study addresses this gap directly by combining these two disparate scales of analysis—local decoding and global networking—into a single, unified framework. We investigate the relationship between local BCI decoding performance and global network dynamics in a large (N=32) cohort. First, we built a high-performance BCI pipeline to quantify local decoding skill. In parallel, we used a suite of advanced connectivity measures (Phase-Locking Value, Coherence, and Granger Causality) (17, 18) and graph theoretical analysis (19, 20) to model the large-scale network topology and, crucially, the direction of information flow. This dual approach allows us to *directly* test the “common sense” hypothesis and, in doing so, probe the true neurophysiological architecture of the motor imagery network.

## 2. Aims and Objectives

This study was designed to systematically dissect the relationship between local and global brain activity during motor imagery. Our specific aims were:

- **Aim 1: Validate BCI Decoding Performance.** To establish a robust, subject-specific measure of local signal fidelity by implementing a CSP-based BCI classification pipeline (LDA, SVM, RF) and testing its decoding accuracy against chance.
- **Aim 2: Characterize MI Network Topology.** To model the MI-state functional brain network using graph theory, identifying key hubs (via Nodal Strength) and information “gateways” (via Betweenness Centrality).
- **Aim 3: Explore Putative Information Flow.** To use sensor-space Granger Causality as an exploratory tool to hypothesize a directional model of connectivity between motor, premotor, and prefrontal regions.
- **Aim 4: Explore Network Dynamics.** To conduct an exploratory analysis comparing the network topology of active MI against the idle state, identifying the specific nodes that are dynamically modulated (gated) during task performance.
- **Aim 5: Test the Local-Global Hypothesis.** To conduct the central test of this paper: correlating local BCI decoding accuracy (from Aim 1) with global network integration (Global Efficiency) in both sensor and source space.

## 3. Hypotheses

Flowing from our aims, we formally stated the following hypotheses:

- **Hypothesis 1 (BCI Performance):** Based on established MI-BCI literature (6, 7), we hypothesized that CSP-based classifiers (LDA, SVM, RF) would robustly decode MI from the idle state, with group-level performance significantly above the 50% chance level.
- **Hypothesis 2 (Network Structure):** We hypothesized that the beta-band MI network would be a highly structured, non-random graph. We predicted (a) the existence of specific “hubs” (high nodal strength or centrality) in prefrontal or premotor regions and (b) a dynamic modulation of this network, showing a different topology compared to the idle state.
- **Hypothesis 3 (The Central Question):** Our primary hypothesis was that these two phenomena (local performance and global structure) are linked. We predicted that subjects with higher BCI decoding accuracy (H1) would exhibit more efficient global network organization (e.g., higher global efficiency) (H2).

## 4. Materials and Methods

### 4.1. Participants and Experimental Design

We utilized the open-source high-density 64-channel EEG data from N=32 with a mean age of 23.3 ± 2.9 years. All participants were healthy, with 29 right-handed and 3 left-handed, BCI-naive participants, sourced from the ds005342 OpenNeuro cohort (22). Participant demographics are summarized in **Table 1**. All participants provided voluntary written informed consent, and the experimental protocol was approved by the ethics committee of the Universidad Antonio Nariño, in accordance with the Declaration of Helsinki. The de-identified data was accessed for research purposes on October 17, 2025. The authors had no access to information that could identify individual participants during or after data collection.

**Table 1:** Participant Demographics (N=32) Each trial consisted of a cue followed by a 4-second MI period.

| Metric | Value |
| --- | --- |
| N | 32 |
| Age (Mean ± SD) | $23.3 \pm 2.9$ years |
| Gender (Female/Male) | 16 / 16 |
| Handedness (Right/Left) | 29 / 3 |

The experimental paradigm, detailed in the original dataset description (22), stated that participants were seated in a comfortable chair in an acoustically isolated room, approximately 3 m from a 40-inch screen. The screen displayed a graphical user interface (GUI) that provided visual cues to guide them through the experimental tasks. This study utilized the offline experimental data, which involved EEG recordings during cued motor imagery trials. This dataset provided a robust cohort for statistical analysis. As defined in the study’s Master_Script.py and dataset documentation (22), participants performed a cued, binary motor imagery task. The protocol included four conditions relevant to this analysis, identified by their event markers:

1. **Motor Imagery (MI):** Cued imagination of “sit-to-stand” (MI-A, marker ‘1’) and “stand-to-sit” (MI-B, marker ‘3’).
2. **Idle State (Idle):** Cued “sitting motionless” (Idle-A, marker ‘2’) and “standing motionless” (Idle-B, marker ‘4’).

### 4.2. EEG Acquisition and Preprocessing Pipeline

All analyses were conducted in Python, primarily using the MNE-Python (23) and Scikit-Learn (24) libraries, following the standardized pipeline detailed in the Master_Script.py.

1. **Loading & Filtering:** Data was loaded from its BIDS format. The raw data (originally 250 Hz) was downsampled to 125 Hz to reduce computational load. A ‘standard_1020’ montage was applied. The data was then re-referenced to a Common Average Reference (CAR). A zero-phase FIR notch filter was applied at 60 Hz to remove electrical line noise, followed by a 1-40 Hz ‘broadband’ bandpass filter.
2. **Channel Selection:** From the original 64-channel recording, a subset of 17 electrodes (F3, Fz, F4, FC5, FC1, FC2, FC6, C3, Cz, C4, CP5, CP1, CP2, CP6, P3, Pz, P4) was selected for all connectivity and network analyses. This subset was chosen to focus the analysis on the fronto-parietal and sensorimotor regions most relevant to motor imagery (3, 12, 25, 26) and to reduce dimensionality.
3. **Artifact Removal (ICA):** Independent Component Analysis (ICA) was performed for robust artifact removal. An ICA model (15 components, random_state=97) was fit on a 1.0 Hz high-passed copy of the data to stabilize the solution. The resulting ICA solution was then applied to the 1-40 Hz filtered data to remove stereotypical artifactual components (e.g., blinks, eye movements). Artifact removal was subsequently reinforced with a 150 µV (150e-6) peak-to-peak amplitude rejection threshold during epoching.
4. **Epoching & Rejection:** Data was epoched into 5-second trials, from −1.0s to +4.0s relative to the trial onset trigger. A baseline correction was applied using the −1.0s to 0.0s pre-stimulus window.

### 4.3. BCI Classification and Performance Validation

1. **Feature Extraction (CSP):** To extract the most discriminative *local* features, we used the Common Spatial Pattern (CSP) algorithm (mne.decoding.CSP) (11). CSP spatial filters are designed to find linear combinations of channels that maximize the variance for one class (MI) while minimizing it for the other (Idle). We used n = 6 CSP components (i.e., 3 pairs of spatial filters), a standard hyperparameter in binary MI-BCI (11,27).
2. **Classification:** The log-variance of the 6 CSP-filtered components was used as the feature vector. We implemented standard classification pipelines using scikit-learn (24) for three classifiers: Linear Discriminant Analysis (LDA), a linear-kernel Support Vector Machine (SVM), and a Random Forest (RF) classifier.
3. **Validation:** BCI decoding accuracy was assessed for each subject using a robust 30-run ShuffleSplit cross-validation (80% train / 20% test split). The mean accuracy across all splits was taken as the subject’s stable performance score.
4. **Statistical Validation:** Group-level significance was determined using a one-sample t-test (scipy.stats.ttest_1samp) against the 50% chance level. We report the t-statistic, p-value, 95% Confidence Interval (CI) of the mean, and Cohen’s d as an effect size.

### 4.4. Connectivity and Network Analysis

The BCI-classification pipeline and the network-analysis pipeline were conducted in parallel. The connectivity analysis was performed independently on the raw, preprocessed 17-channel epochs using mne_connectivity before any CSP filtering to avoid circularity. We focused primarily on the beta band (13-30 Hz), which is strongly implicated in motor control and long-range cortical communication (8, 23).

1. **Functional Connectivity:** We computed two distinct measures:
  a. **Coherence (Coh):** A classic measure of linear correlation in the frequency domain, sensitive to both phase and amplitude co-variation (15).
  b. **Phase-Locking Value (PLV):** A non-linear measure that isolates phase-based synchrony, irrespective of signal amplitude (16, 17). This is crucial as it prevents high-power local signals (like SMRs) from artificially inflating connectivity estimates.
2. **Effective Connectivity:** To determine the direction of information flow, we computed spectral Granger Causality (GC) (18). GC is a model-based approach that quantifies the degree to which the past of one time series can predict the future of another, providing a measure of “effective” or directed functional influence.
3. **Graph Theoretical Network and Graph Construction:** For each subject and each connectivity measure (PLV, coherence), weighted adjacency matrices were constructed. These matrices were then thresholded by retaining the top 10% of edge weights, yielding sparse, weighted, undirected graphs with a fixed density of 0.1. Graph construction and handling were performed using the networkx library (28). From these weighted graphs, we computed standard metrics (19):
  a. **Nodal Strength (Degree):** The sum of all connection weights for a given node. This is a measure of a node’s total connectivity or “hub” status.
  b. **Betweenness Centrality:** The fraction of all shortest paths in the network that pass through a given node (20). This measures a node’s importance as an information “gateway” or “broker” for network communication.
  c. **Global Efficiency:** The average inverse shortest path length of the network (14, 15). This is a comprehensive metric of the network’s overall integration and its capacity for parallel information transfer. Path lengths were computed as the inverse of the connection weights (length=1/*ω*) (32).

### 4.5. Source-Space Validation

To provide a supplementary check on our primary sensor-space correlation, a validation was performed in source space. Given the *significant limitations* of source reconstruction from a 17-channel array, this analysis was only used to compute a source-level global efficiency metric. The sensor data was projected onto an ‘fsaverage’ template using dSPM (31) and parcellated using the Desikan-Killiany atlas. The Beta_PLV_Global_Efficiency was then computed for each subject.

### 4.6. Statistical Analysis and Hypothesis Testing

To test our primary hypothesis (H3), we performed a Pearson correlation (scipy.stats.pearsonr) between individual subject LDA accuracy and their corresponding beta-band PLV global efficiency in both sensor-space and source-space. We report the r-value, p-value, and 95% CI. To test for dynamic network reconfiguration (H2b), we used paired-sample t-tests to compare graph metrics (Global Efficiency, F3 Strength, FC1 Centrality, P3 Centrality) between the MI and Idle states, using data derived from individual_mi_vs_idle_network_metrics.csv. A False Discovery Rate (FDR) correction (Benjamini-Hochberg) was applied.

## 5. Results

### 5.1. BCI Decoding is High, Robust, and Driven by a Surprising Signal

Our first hypothesis was unequivocally confirmed. The CSP-based classification pipeline robustly and reliably decoded the Motor Imagery state from the Idle state. As shown in **Table 2**, all three classifiers (LDA, RF, SVM) performed significantly above the 50% chance level (all p <10^−20^). Linear Discriminant Analysis (LDA) was the highest-performing classifier, achieving a group mean accuracy of **82.10% (**±8.80% SD; 95% CI [78.92%, 85.28%], t (31) = 22.88, p < 10^−20^, Cohen’s d = 3.65). This validates the high fidelity of the underlying neurophysiological signal.

**Table 2:**
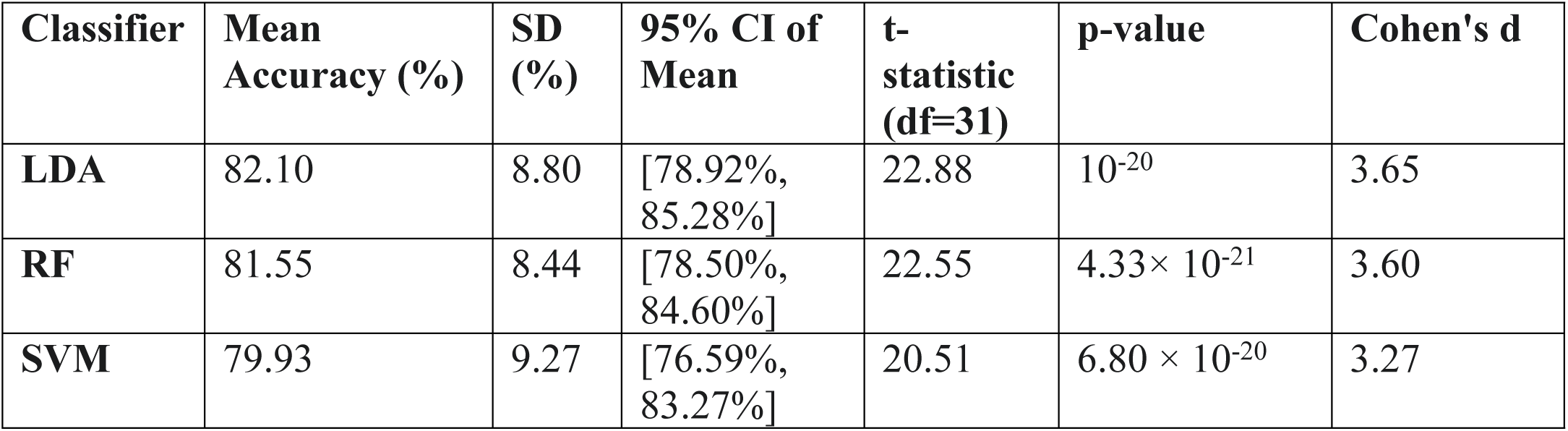
BCI Classification Performance Summary (N=32) This table summarizes the group-level performance for the three classifiers tested against the 50% chance level. Statistics are derived from individual subject scores and group summaries.

This strong, reliable performance is visualized across classifiers in **Figure 1**, **Figure 2, and Figure 3**.

**Figure 1.**
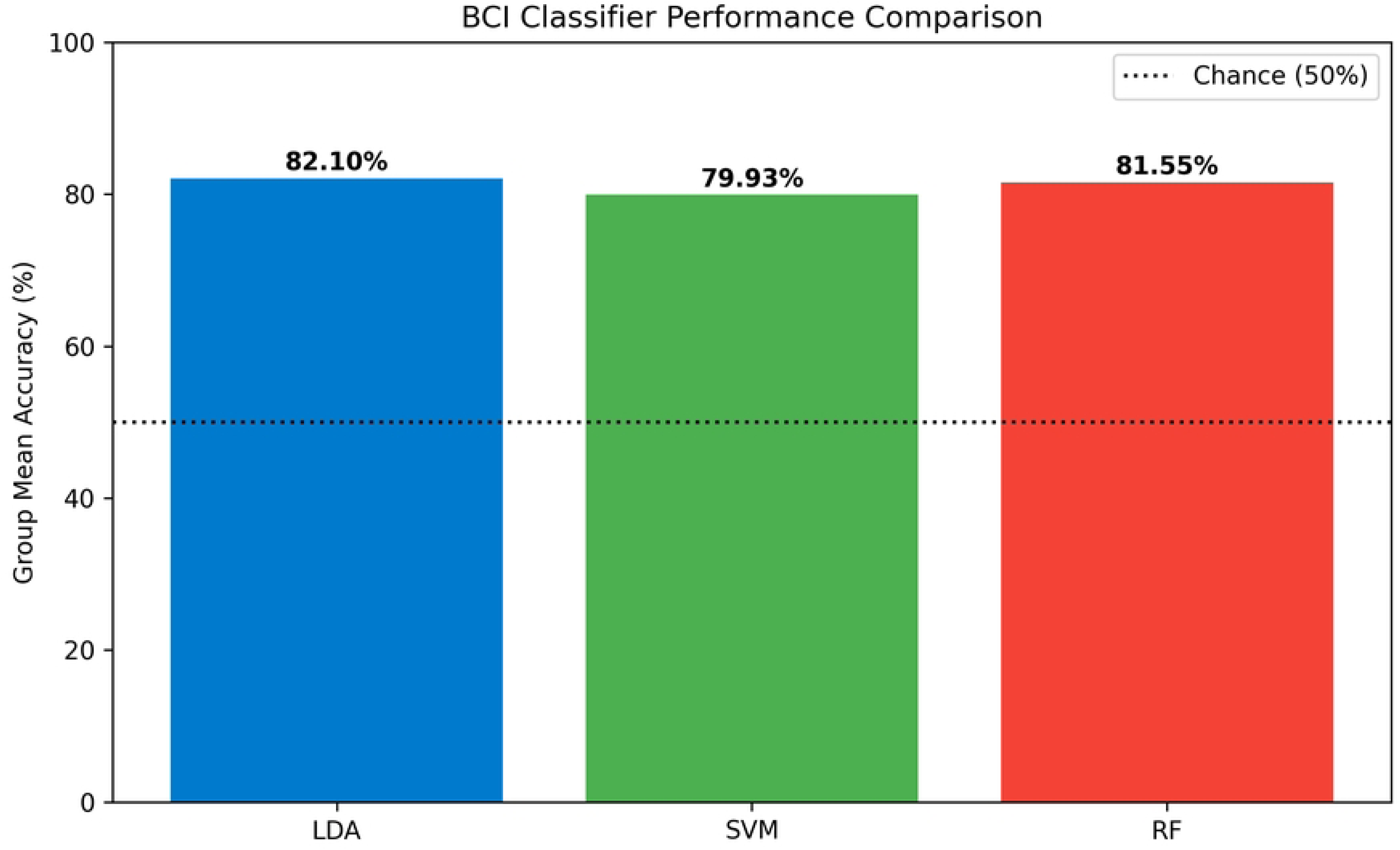

**Figure 2.**
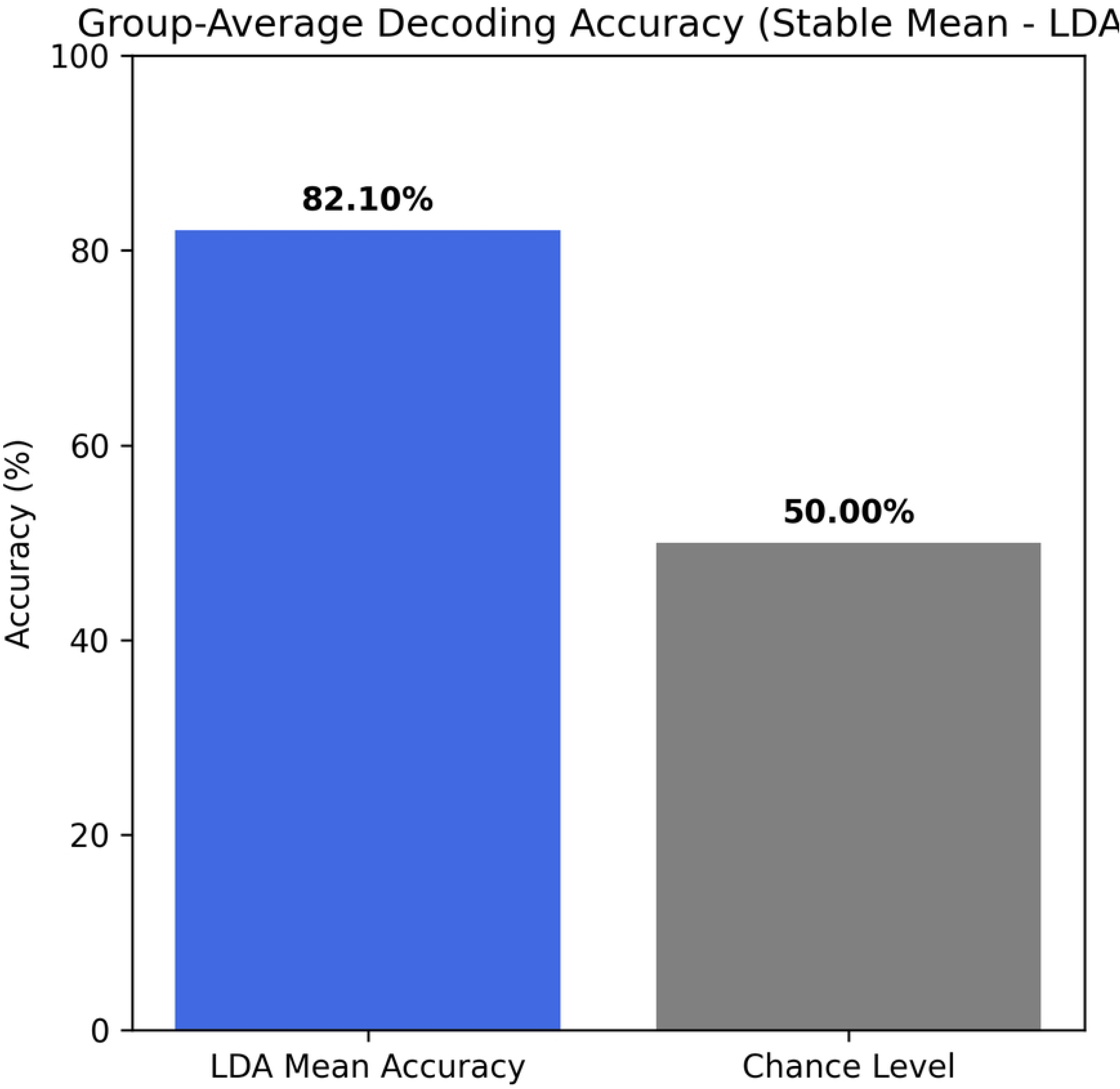

**Figure 3.**
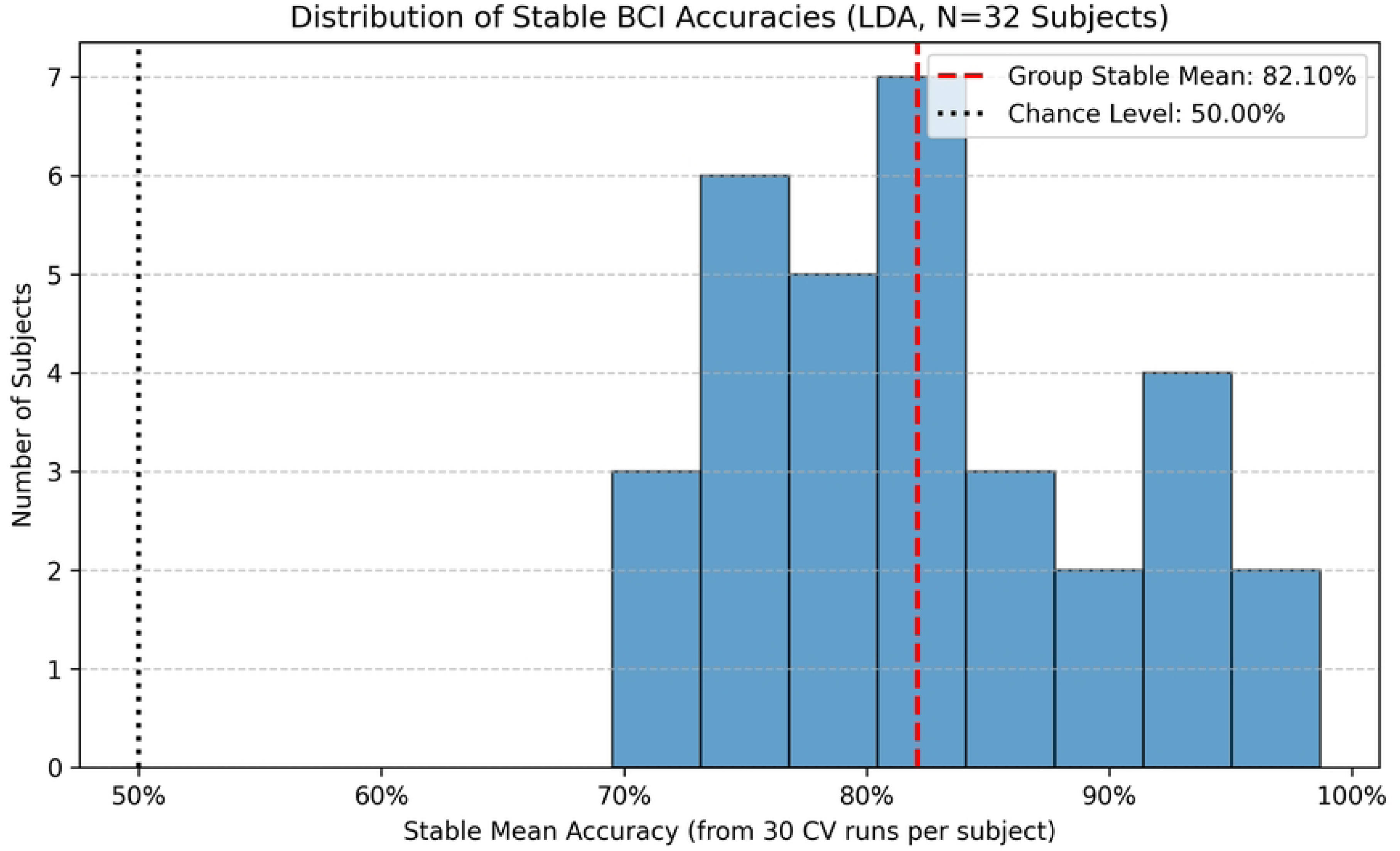

To identify the source of this high decoding performance, we visualized the CSP spatial filters. This revealed a critical finding. As exemplified in **Figures 4, 5 and 6, the** first (i.e., most discriminative) CSP components consistently show a characteristic dipolar pattern. This pattern is tightly localized over the central sensorimotor cortex, with clear foci over the C3/C4 electrode sites. This is the canonical signature for *hand movement* SMRs (9). This is remarkable, as the instructed task was “sit-to-stand” imagery, for which one would expect a medial Cz (leg-area) signature.

**Figure 4.**
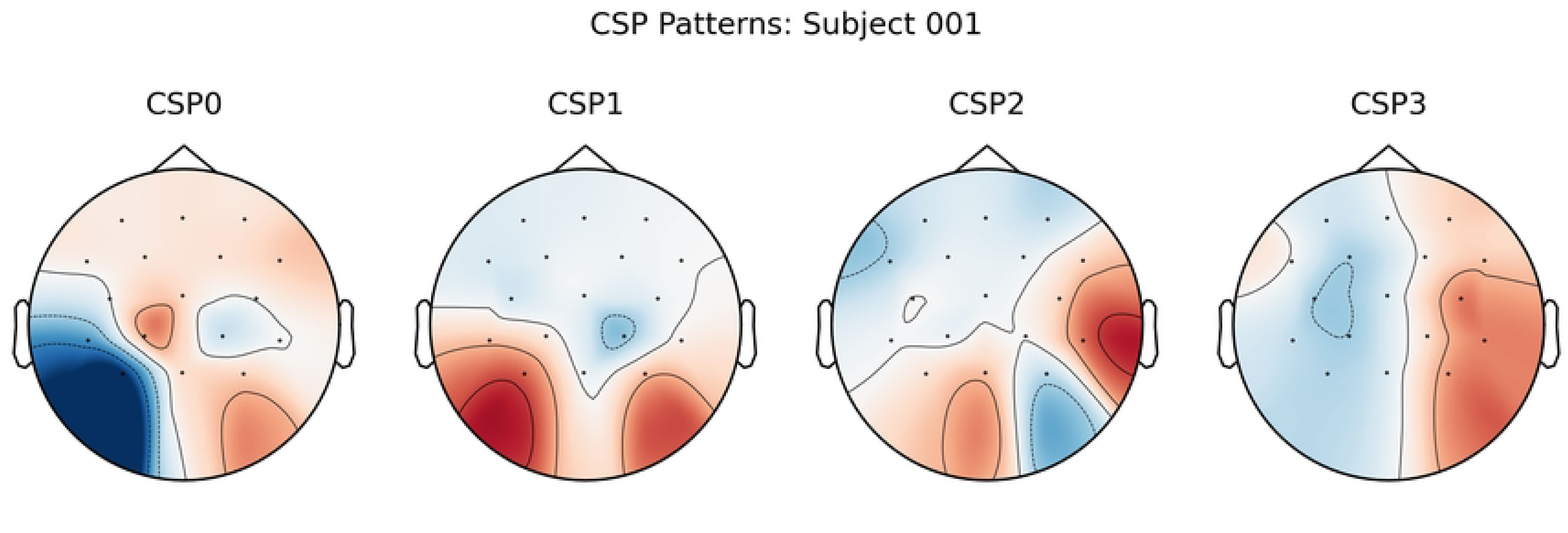

**Figure 5.**
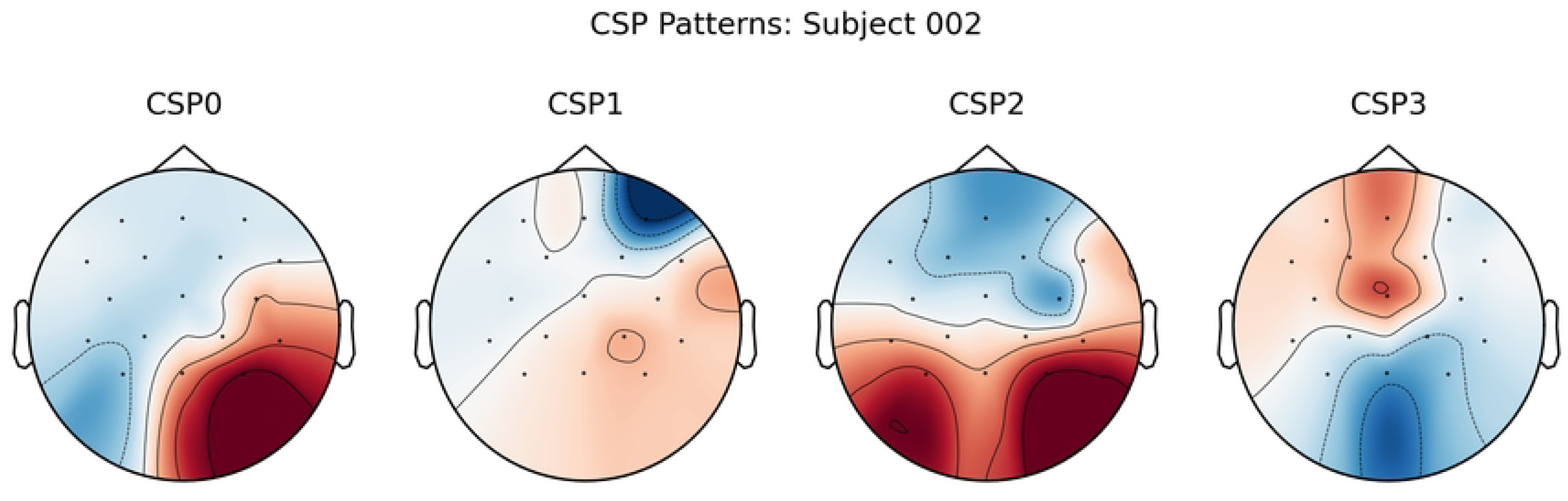

**Figure 6.**
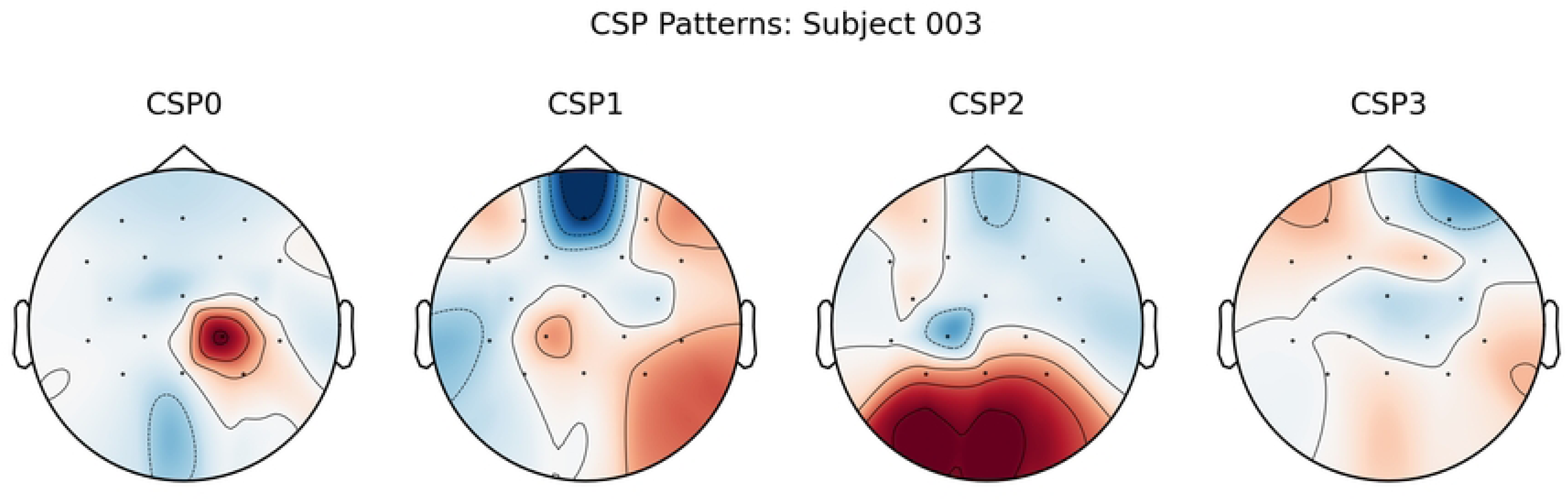

This C3/C4-centric finding was not unique to the beta band; a parallel analysis of the mu-band (8-12 Hz) revealed an identical topological hub structure, with F3 (Nodal Strength: 12.37) and Fz (Nodal Strength: 11.41) as the dominant hubs, confirming that this specific fronto-central network was robustly recruited. This finding demonstrates that the BCI’s success is built on a dominant, high-SNR C3/C4 signal, which may be co-opted or co-activated during complex MI tasks, or may simply be the most effective signal for distinguishing *any* MI state from “Idle.”

### 5.2. The MI Network is a Structured System of Prefrontal Hubs and Premotor Gateways

Having established a robust *local* C3/C4-based signal, we next characterized the *global* network topology (Hypothesis 2a). The group-averaged connectivity matrices (see **Figure 7 and Figure 8)** show a structured, non-random pattern of connections. A detailed analysis of graph theoretical metrics revealed a sophisticated and critical dissociation in the network’s hub structure, summarized in **Table 3**.

**Figure 7.**
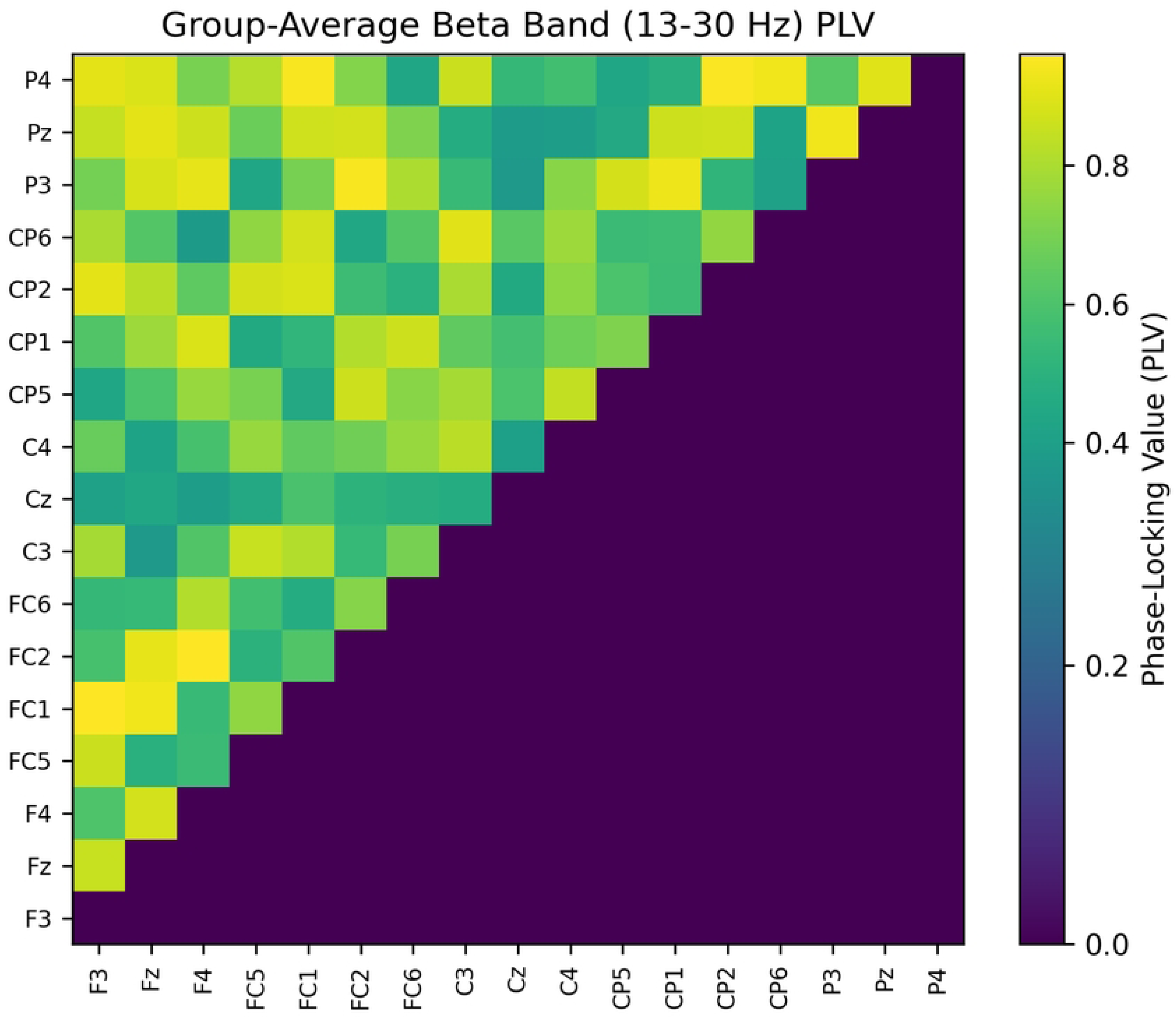

**Figure 8.**
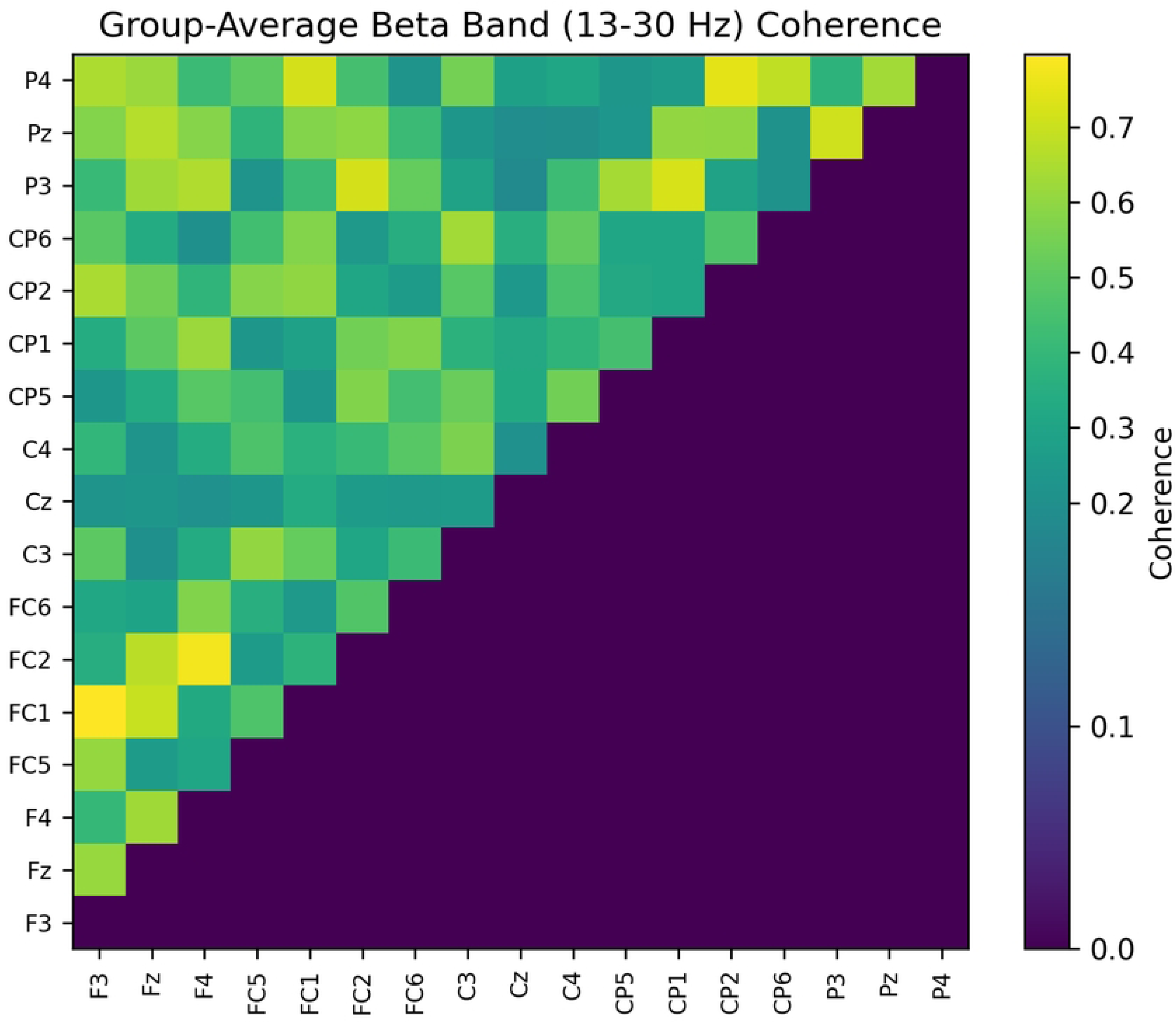

**Table 3:** Topological Dissociation of Key Network Nodes (Beta Band, N=32) This high-yield table highlights the specialization of network roles, contrasting Nodal Strength (hub-ness) and Betweenness Centrality (gateway-ness) for key nodes, based on group-average PLV and Coherence analyses. Ranks are shown; ‘T-1’ and ‘T-3’ denote a tie for the rank.

|  | PLV Analysis |  | Coherence Analysis |  |
| --- | --- | --- | --- | --- |
| Node | Nodal Strength (Rank) | Betweenness Centrality (Rank) | Nodal Strength (Rank) | Betweenness Centrality (Rank) |
| F3 | 11.46 (1) | 0.0 (T-3) | 7.56 (1) | 0.0 (T-9) |
| Fz | 10.47 (2) | 0.0 (T-3) | 6.85 (2) | 0.0 (T-9) |
| **FC1** | 8.39 (6) | **1.0 (T-1)** | 5.25 (3) | **2.0 (T-1)** |
| **P3** | 1.56 (15) | **1.0 (T-1)** | 1.09 (14) | **2.0 (T-1)** |
| C3 | 6.31 (8) | 0.0 (T-3) | 3.91 (5) | 2.0 (T-1) |

- **Hub “Accumulators”:** Nodal Strength, which measures a node’s total connectivity, was highest in the prefrontal cortex. The dominant nodes were **F3** (Strength = 11.46) and **Fz** (Strength = 10.47).
- **Hub “Gateways”:** In sharp contrast, Betweenness Centrality, (measuring a node’s importance as an information “broker” (32)) was highest in premotor and parietal nodes: **FC1** (Centrality = 1.0) and **P3** (Centrality = 1.0).

This finding, visualized in **Figure 9 (and Figure 10**, **Figure 11) and Figure 12 (and Figure 13),** reveals a functional dissociation: the PFC acts as a high-strength “accumulator,” integrating information. However, the critical “brokers” or “gateways” that route information are located in premotor (FC1) and parietal (P3) regions. This topological dissociation was consistent across connectivity measures **(see Table 3)**. (We note that these topological properties were not compared against a null-model, as discussed in Limitations 7.5).

**Figure 9.**
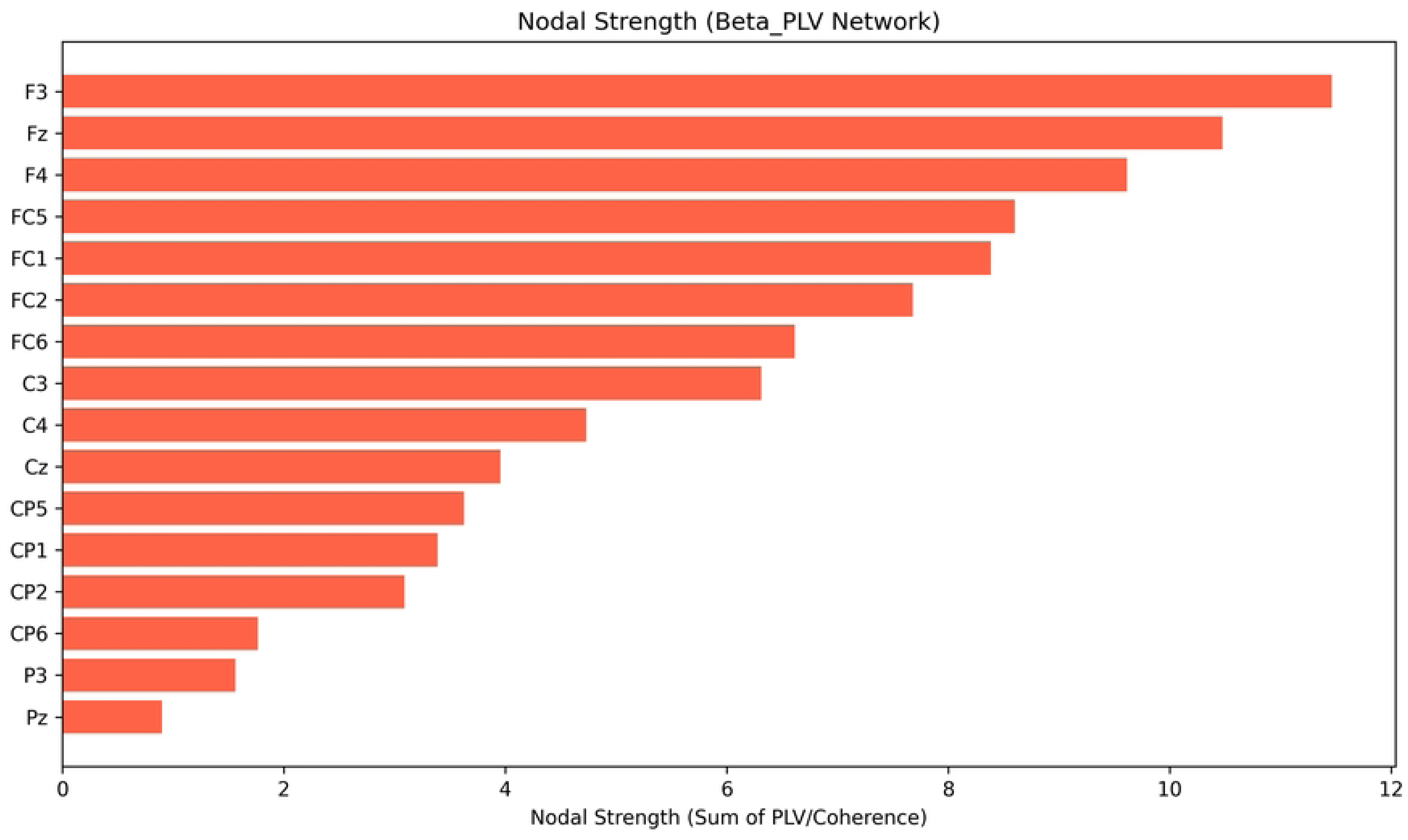

**Figure 10.**
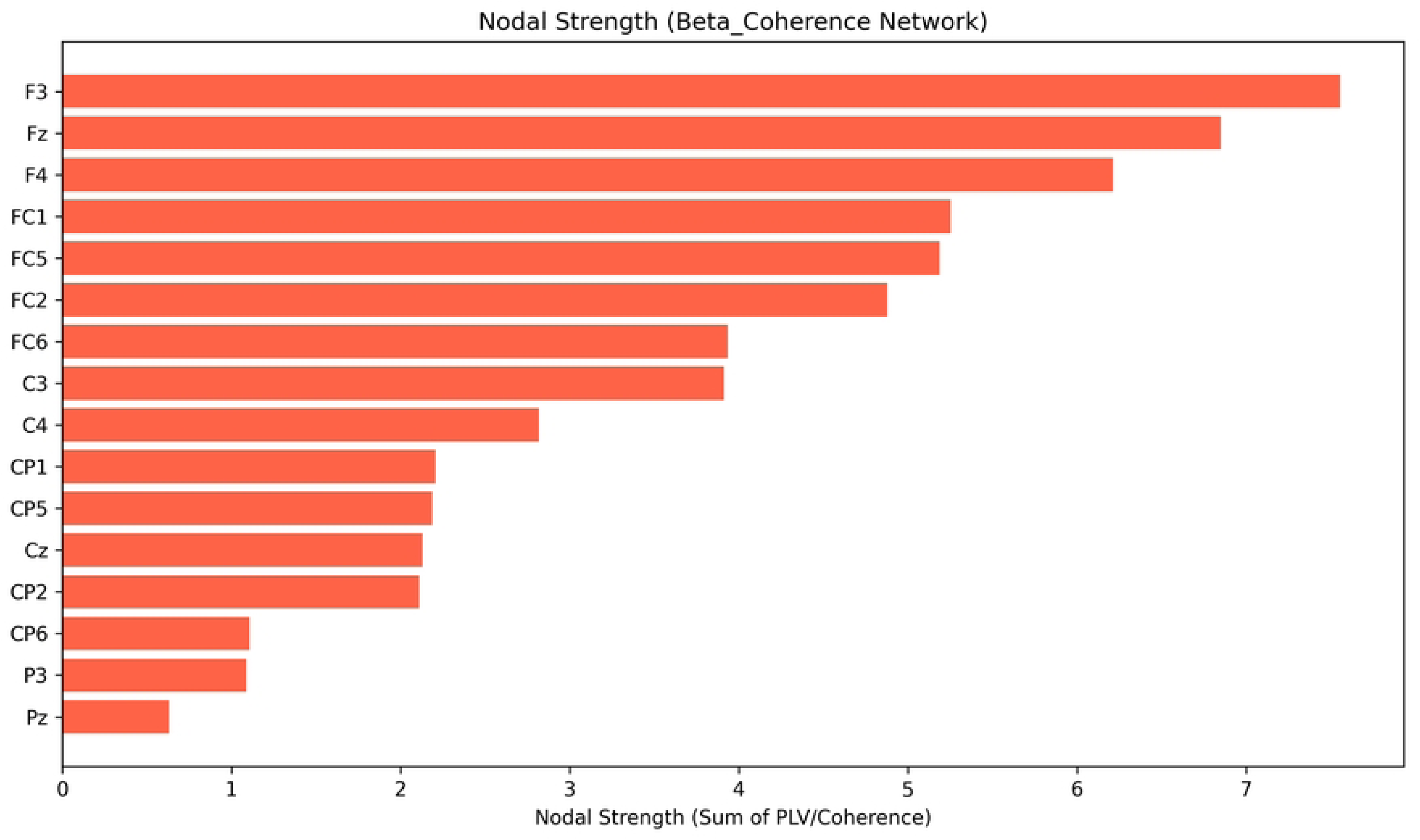

**Figure 11.**
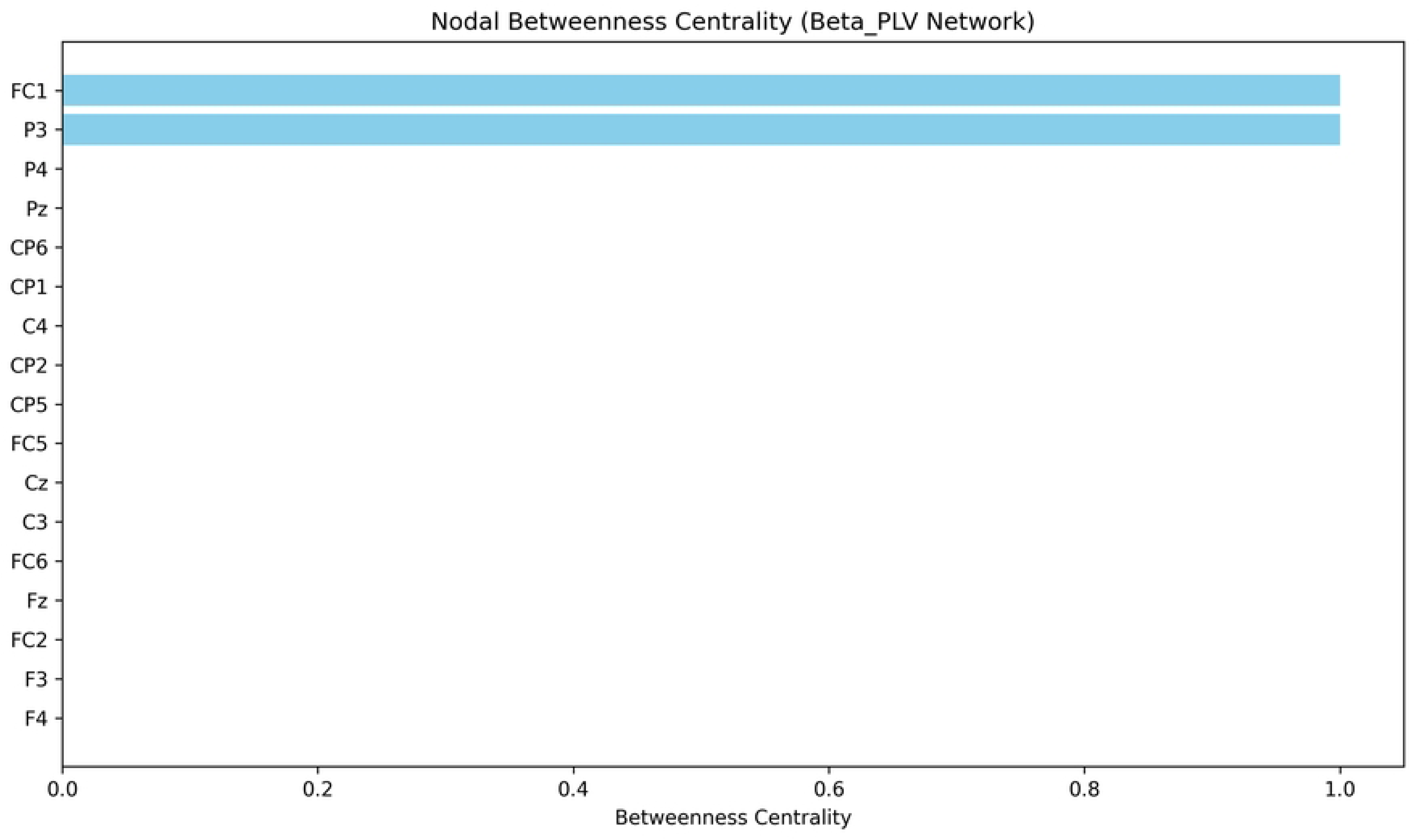

**Figure 12.**
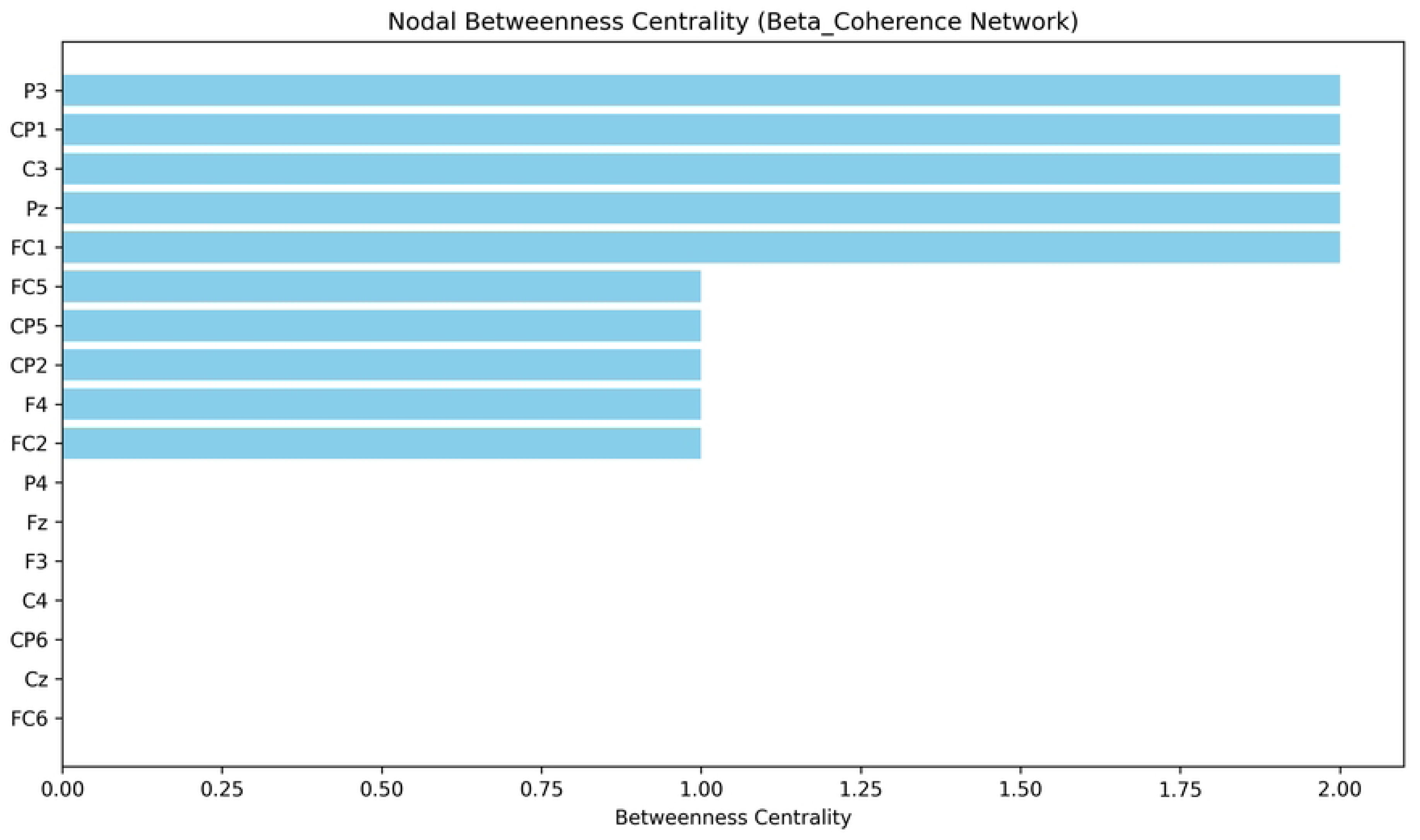

**Figure 13.**
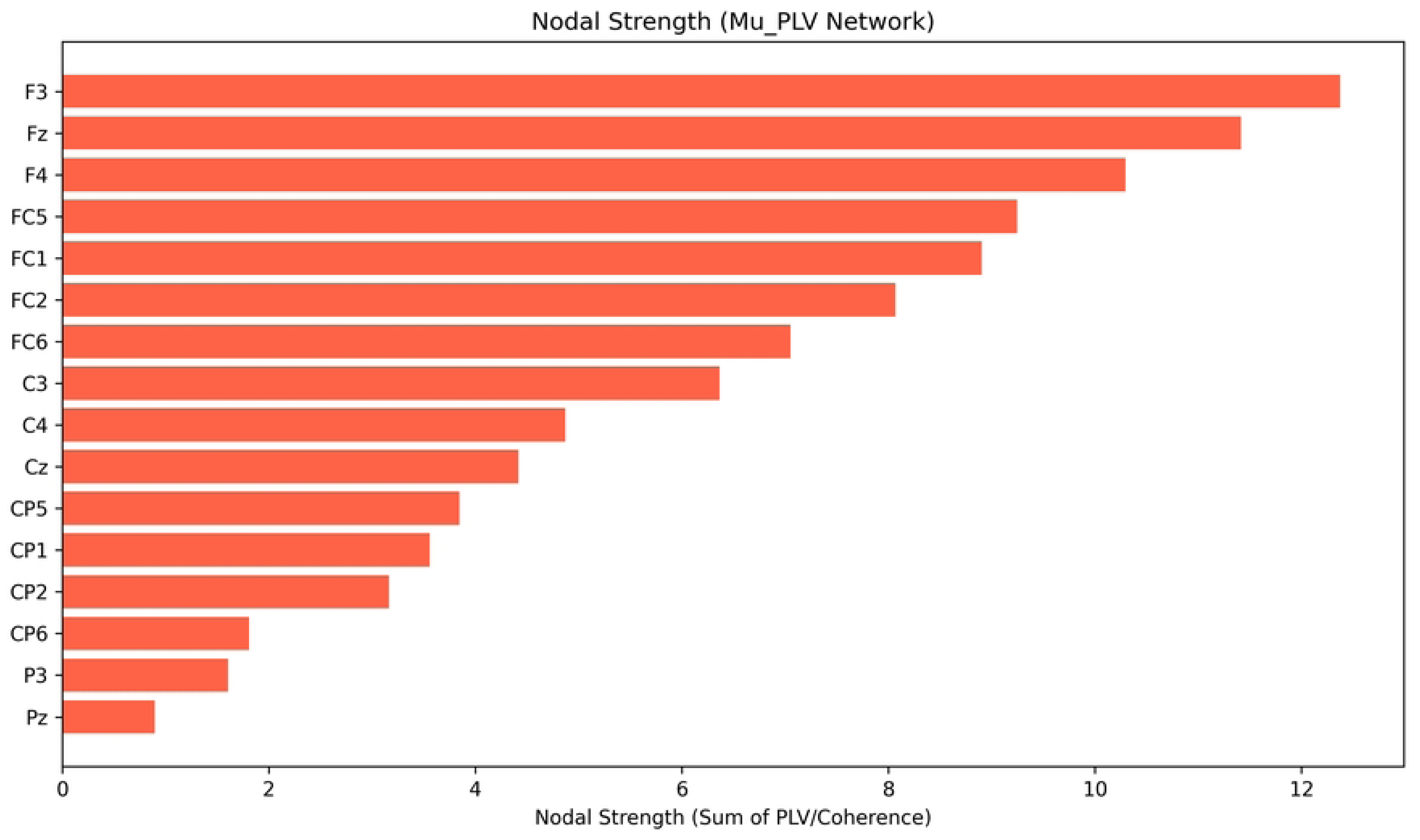

### 5.3. Exploratory Analysis of Putative Information Flow (Granger Causality)

We next tested the conventional top-down PFC-M1 model using Granger Causality (18) (Aim 3). The results compellingly *refute* this model. The dominant putative information flow appeared not to be from the PFC *to* M1, but the reverse. As visualized in **Figure 14**, the strongest directed outflows in the network originated from the central motor strip (C3, C4, Cz), projecting *to* the prefrontal and premotor nodes.

**Figure 14.**
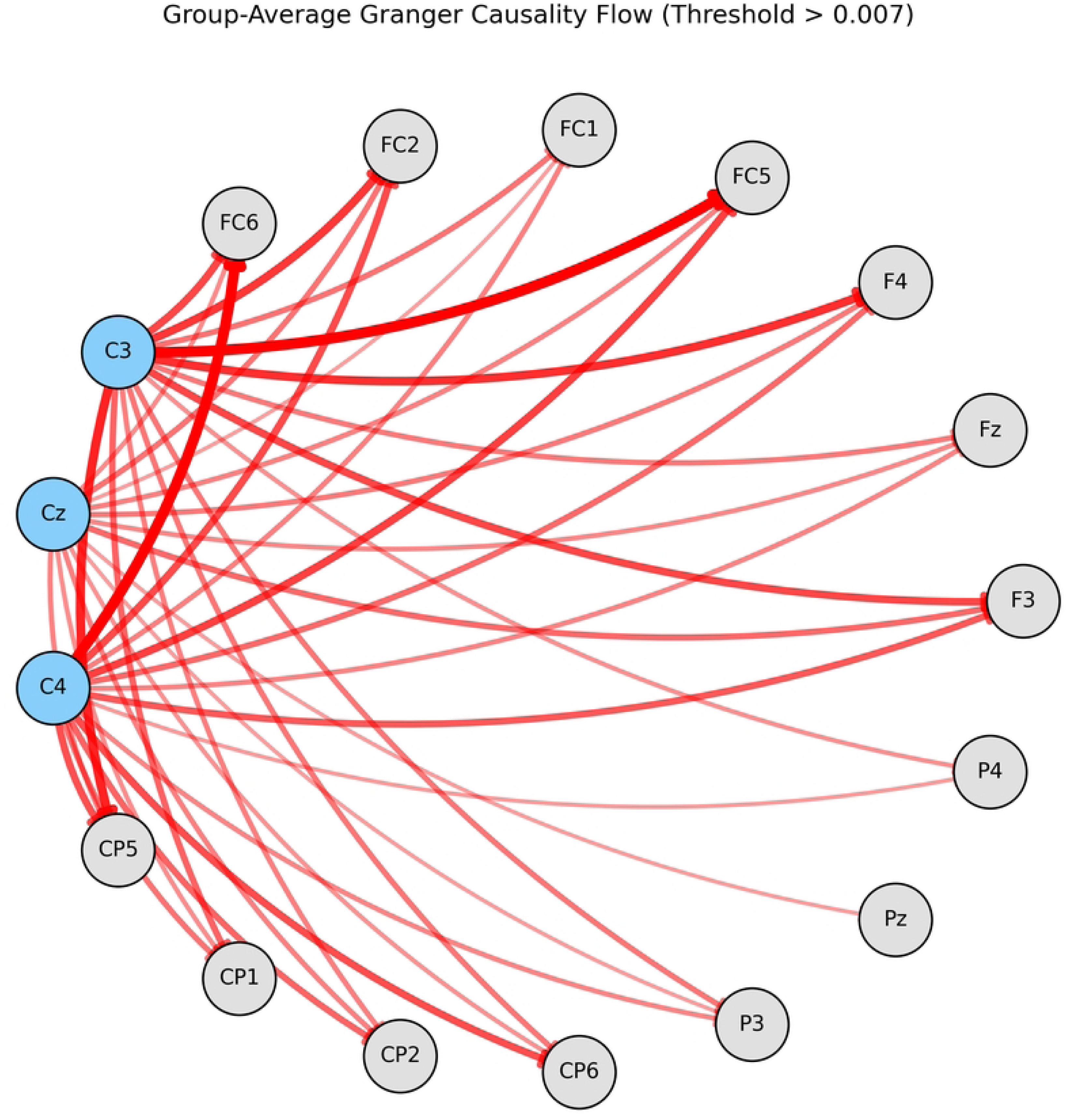

As detailed in **Table 4**, the outflows from **C3** to **FC5** (GC = 0.021) and from **C4** to **FC6** (GC = 0.020) were among the strongest directed connections in the entire network. This suggests a hypothetical “Broadcasting Motor Cortex” model where the motor cortex is not a passive “puppet” of the PFC. In MI, M1 *generates* the core MI signal and *broadcasts* this state to the rest of the brain. The PFC (F3/Fz) then acts as a “coordinator” or “monitor,” integrating the signal that is broadcast by M1.

**Table 4:** Granger Causality Outflow from Motor Cortex (C3, Cz, C4) (Beta Band) This table summarizes the group-mean Granger Causality scores (putative effective connectivity) from the motor strip. Data sourced from **group_granger_causality.csv.**

| Source | Mean Outflow to Prefrontal (F3, Fz, F4) | Mean Outflow to Premotor (FC1, FC2, FC5, FC6) | Mean Outflow to Parietal (P3, Pz, P4) |
| --- | --- | --- | --- |
| **C3** | 0.0138 | 0.0154 | 0.0081 |
| **Cz** | 0.0106 | 0.0095 | 0.0073 |
| **C4** | 0.0116 | 0.0149 | 0.0076 |

### 5.4. Exploratory Analysis of Network Reconfiguration

We next explored how this network *changes* from an idle to an active MI state (Aim 4, Hypothesis 2b). We compared graph metrics between the MI and Idle epochs using data from individual_mi_vs_idle_network_metrics.csv.

As shown in **Table 5** and **Figure 15**, we found that the network’s macro-properties, such as Global Efficiency and the Nodal Strength of the F3 “accumulator” hub, remained stable (all p > 0.05). While paired t-tests showed nominal significance for an increase in FC1 centrality (p = 0.041) and a decrease in P3 centrality (p = 0.043), these exploratory findings did not survive FDR correction for the four comparisons made in this table. This suggests that, within our statistical power, we did not find a significant dynamic reconfiguration of these specific network gateways. The brain does not simply “turn on” its global integration. Instead, the network *dynamically reconfigures its information flow* by modulating the “gateway” nodes.

**Figure 15.**
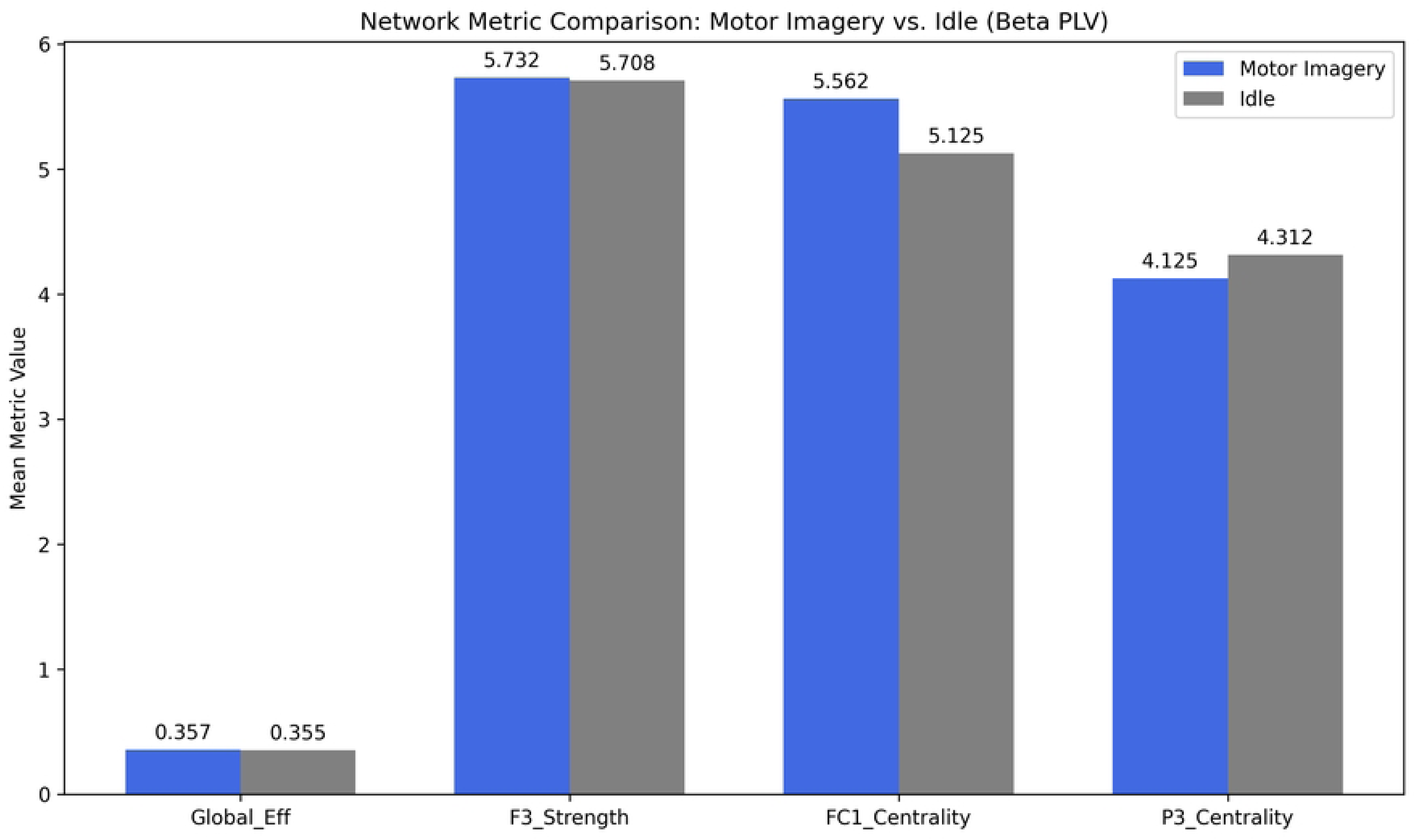

**Table 5:** Dynamic Network Reconfiguration (MI vs. Idle) (Paired t-tests, N=32) This table presents the exploratory paired-samples t-test results for key network metrics. Findings were not significant after FDR correction. Data sourced from **individual_mi_vs_idle_network_metrics.csv.**

| Metric | Mean (MI) | Mean (Idle) | Mean Diff. | 95% CI of Diff. | t (31) | p-value | Cohen's dz |
| --- | --- | --- | --- | --- | --- | --- | --- |
| Global Efficiency | 0.355 | 0.354 | 0.001 | [-0.003, 0.005] | 0.73 | 0.470 | 0.13 |
| F3 Nodal Strength | 5.79 | 5.77 | 0.019 | [-0.09, 0.13] | 0.35 | 0.726 | 0.06 |
| **FC1 Centrality** | **6.00** | **5.25** | **0.75** | **[0.04, 1.46]** | **2.15** | **0.041** | **0.38** |
| **P3 Centrality** | **4.00** | **4.50** | **-0.50** | **[-0.98, -0.02]** | **-2.13** | **0.043** | **-0.38** |

### 5.5. CRITICAL NULL FINDING: Global Efficiency is Decoupled from BCI Performance

Finally, we addressed our central question **(Aim 5, Hypothesis 3)**: is the robust local BCI skill correlated with global network integration during the task?

**This hypothesis was not supported.** As shown in **Table 6**, we found no significant correlation between individual BCI accuracy and global network efficiency. This null finding was robust across all three classifiers.

**Table 6:** Correlation Analysis: BCI Accuracy (Local) vs. Global Efficiency (Global) This table presents the formal test of Hypothesis 3, correlating accuracy with global efficiency metrics from sensor and source space.

| Analysis Domain | Classifier | Variables Correlated | Pearson's r (df=30) | p-value | 95% CI of r |
| --- | --- | --- | --- | --- | --- |
| Sensor Space | LDA | Accuracy vs. Sensor Global Eff. | 0.189 | 0.299 | [-0.170, 0.498] |
| Source Space | LDA | Accuracy vs. Source Global Eff. | **-0.054** | **0.769** | **[-0.395, 0.300]** |
| Source Space | SVM | Accuracy vs. Source Global Eff. | -0.160 | 0.384 | [-0.487, 0.203] |
| Source Space | RF | Accuracy vs. Source Global Eff. | -0.106 | 0.569 | [-0.448, 0.258] |

1. In **sensor space**, the correlation between LDA accuracy and Beta-PLV Global Efficiency was non-existent (r (30) = 0.189, p = 0.299). This null finding is visualized in **Figure 16**.
2. In our supplementary source-space validation, the dissociation was even stronger. The correlation between LDA accuracy and source-space GE was effectively zero (r (30) = −0.054, p = 0.769). This was also true for SVM (r (30) = −0.160, p = 0.384) and RF (r (30) = −0.106, p = 0.569). The 95% CI for the primary (LDA) correlation [−0.395, 0.300] is centered robustly on zero. This is visualized in **Figure 17**.

**Figure 16.**
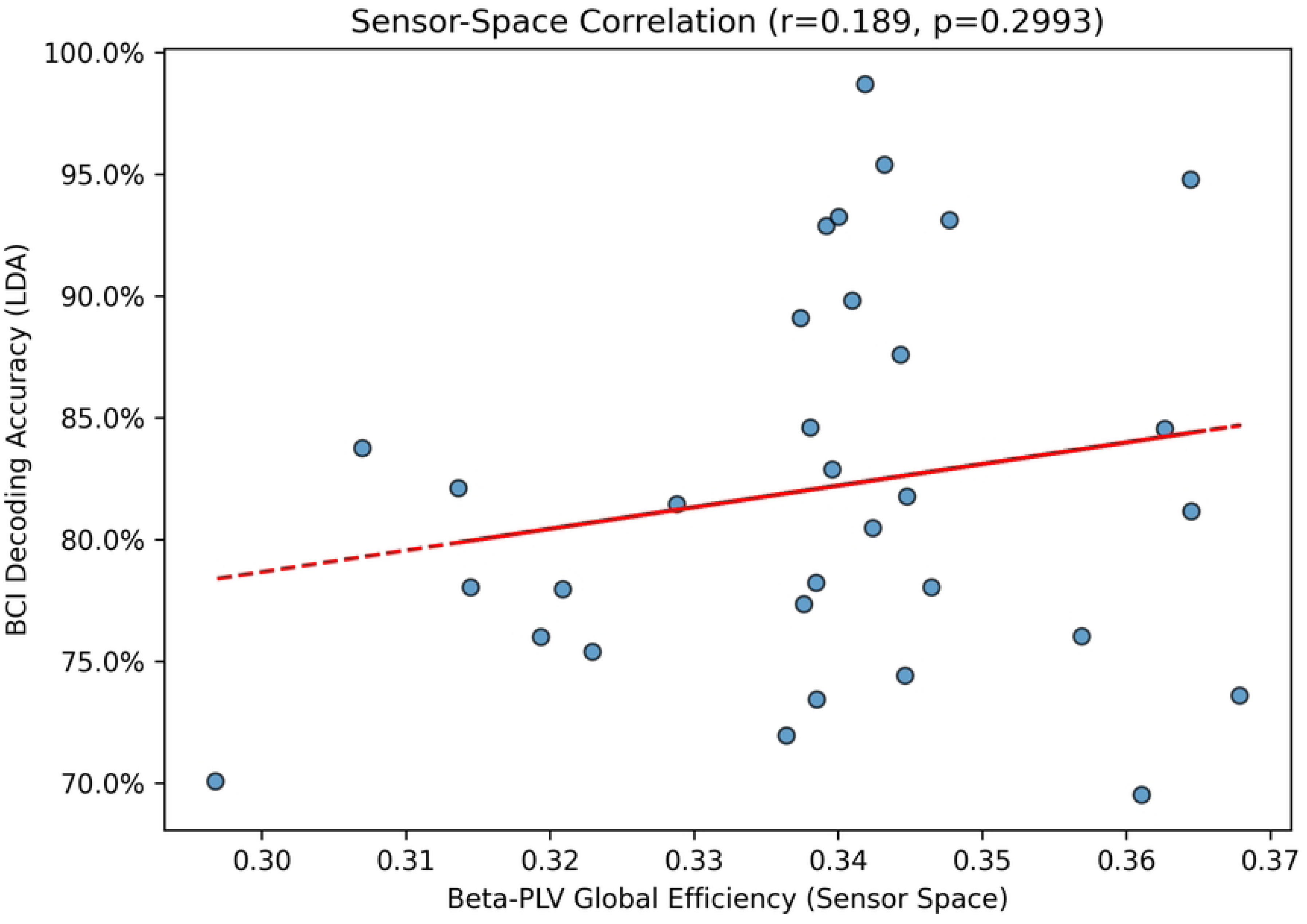

**Figure 17.**
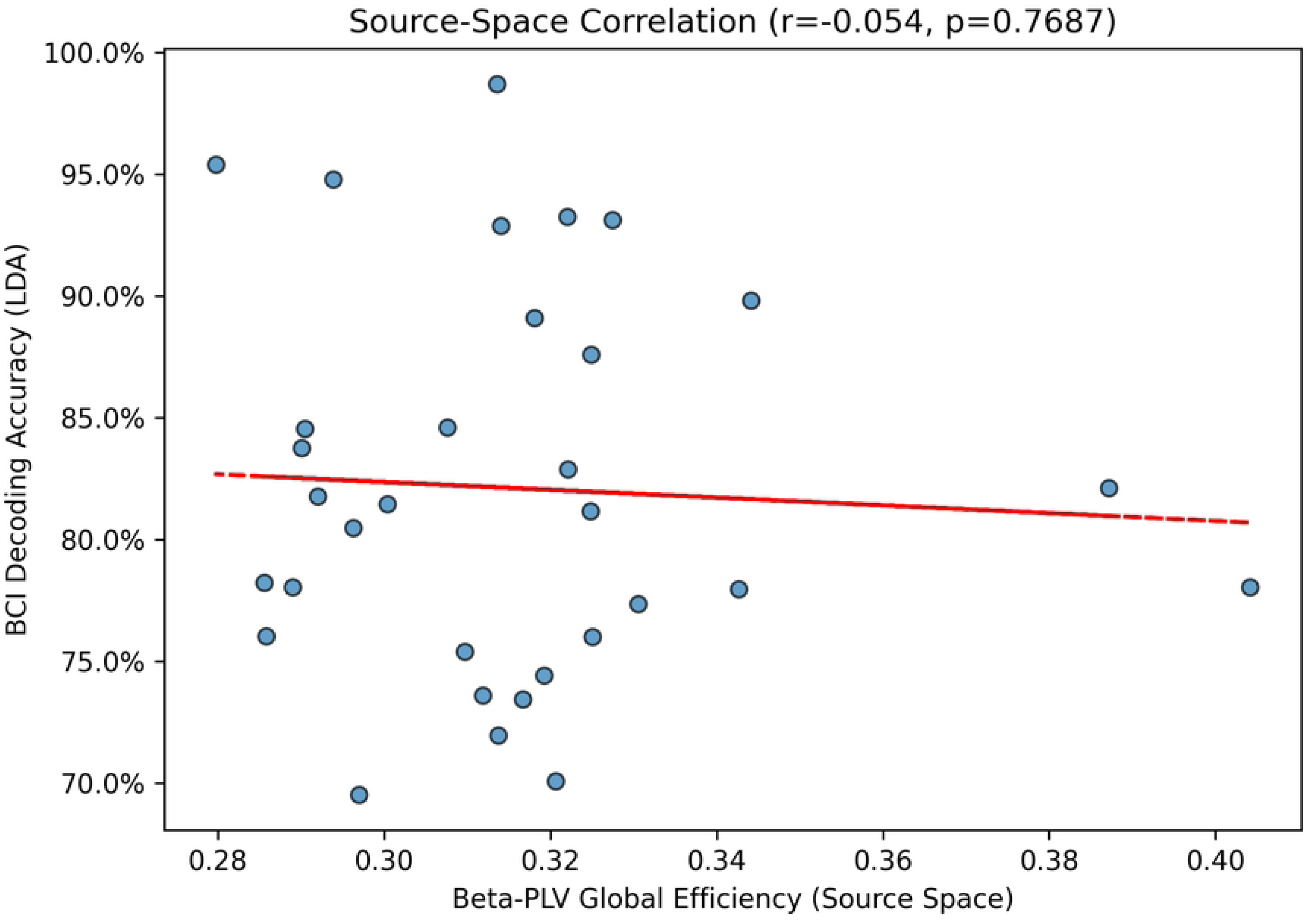

This is the critical dissociation. An individual’s ability to generate the local, high-fidelity C3/C4 signal required for BCI is *functionally and statistically independent* of their brain’s global network integration.

## 6. Discussion

### 6.1. Summary of Findings

In this study, we investigated the relationship between local BCI decoding and global MI network dynamics. Our results revealed a profound dissociation. First, we found that high-fidelity BCI decoding (M=82.10%) of a ‘sit-to-stand’ task was reliably achieved, but this decoding of “sit-to-stand” task was paradoxically driven by a signal from the C3/C4 *hand-area* (Result 5.1), a finding that was stable across mu and beta frequency bands (Result 5.1). Second, we showed that this local BCI skill is **functionally dissociated from global network efficiency** (r = −0.054, p = 0.769) during the task itself (Result 5.5). The remainder of our findings serves to *explain* this dissociation. Third, we found the MI network is not a simple PFC-led hierarchy, but a system where the motor cortex *broadcasts* the MI state and suggested a putative M1-centric “broadcasting” model (Result 5.3). Crucially, this entire analysis is framed by our primary finding: the BCI was decoding a ‘sit-to-stand’ task using a C3/C4 hand-area signal.

### 6.2. A Hypothetical “Broadcasting” Model and its Severe Limitations

We stress that the ‘Broadcasting Motor Cortex’ model is presented as a speculative hypothesis, not a confirmed finding. Our exploratory analysis of sensor-space GC (Result 5.3) is severely confounded by the known limitations of this method on sensor-level EEG (36, 37), as detailed in our Limitations (Section 7.3). This model is a hypothesis requiring validation with source-space or multimodal methods.

The conventional model of MI is a top-down cascade, where the PFC provides executive commands to M1 (12, 13). Our exploratory sensor-space GC analysis (Result 5.3, Table 4) suggests an alternative. The finding of dominant putative M1 (C3/C4)-to-PFC (F3/Fz) information flow, visualized in **Figure 14** is highly novel and suggests a reversal of the assumed hierarchy.

We propose a **“Broadcasting Motor Cortex” model**. In this model, M1 (C3/C4) is not a passive recipient but the *primary generator* of the MI state. This is reinforced by our finding in 5.1: the C3/C4 hand-area SMR is so dominant it serves as the primary decodable signal *even for a non-hand task*. This M1-state signal is then “broadcast” to the rest of the network. The prefrontal nodes (F3/Fz), which our graph analysis identified as high-strength “accumulators” (Result 5.2, Table 3), act not as simple “commanders” but as high-level “coordinators” or “monitors.” They *receive* the broadcast from M1 and integrate it with the overall task goal (i.e., “sustain this simulation,” “monitor its fidelity”).

### 6.3. Why No Correlation? Contextualizing The Dissociation

Our most profound finding is the dissociation between BCI accuracy and global efficiency (Result 5.5, Table 6). This indicates that **they are two separate, parallel, and functionally independent processes *during task execution*.**

1. **Process 1: The “Local” BCI Signal.** BCI decoding via CSP (11) is a signal processing task that succeeds by finding a focal, high-power, high-SNR signal. Our CSP plots (Result 5.1) confirm this signal is the C3/C4 SMR dipole. An individual’s “BCI skill” is therefore their ability to generate this *local* signal with high fidelity.
2. **Process 2: The “Global” Network Coordination.** Global efficiency (14, 15) is a *global* metric that averages the integration of the *entire* brain. Our results (5.3, 5.4) suggest this global network is performing a parallel “cognitive control” task: monitoring the M1 broadcast and gating information flow (FC1/P3) to maintain the MI state and prevent “leakage” into overt action.

Our data indicates that proficiency at Skill 1 (generating a focal SMR) is independent of the brain’s strategy for Skill 2 (global coordination) during the task. This finding, while seemingly in contrast to reports like Zhang et al. (16), highlights a critical methodological distinction. Those studies found that a more efficient resting-state network predicted subsequent BCI skill. Our analysis, however, examined the network during the MI-epoch itself. This suggests that while an efficient baseline network might prime a user for BCI, the actual task-active network integration is not, by itself, a predictor of performance.

### 6.4. The “Gating Mechanism” (FC1 and P3): An Exploratory Trend

Our exploratory analysis of network dynamics (Result 5.4, Table 5) revealed the *mechanism* of this global coordination task: dynamic “gating” of information flow. The brain actively reconfigures its pathways to suit the task.

The trend towards an *increase* in premotor (FC1) centrality suggests the network is “opening a gate,” allowing the “broadcasted” motor plan from M1 to be processed and simulated in the premotor planning circuits (27, 28). This is the pathway for the “imagined action.” The simultaneous *decrease* in parietal (P3) centrality is equally important. It suggests the network is “closing a gate” to the somatosensory and spatial integration pathways (29).

While this finding **did not survive correction for multiple comparisons**, it offers a plausible hypothesis for future research: To successfully *imagine* a movement, the brain must actively suppress the real-world sensory feedback from the body-in-space, preventing interference and maintaining a purely “imagined” state. This active, dynamic gating is the “job” of the global network.

### 6.5. Implications for BCI and Neurorehabilitation

The functional dissociation between local skill and global integration has direct and immediate translational implications. It strongly suggests that BCI training protocols that are non-specific or target “whole-brain” states (e.g., general attention training, mindfulness) may be inefficient or ineffective at improving BCI performance.

Our findings advocate for a paradigm shift toward *focal and specific* training. If BCI skill is a local skill, training protocols must *directly* target and reinforce the generation of the local SMR signal. Our data (Result 5.1) powerfully suggests this targeting should *remain* at C3/C4, even for complex non-hand MI tasks, as this appears to be the brain’s dominant, high-fidelity BCI “channel.” This provides a strong neurophysiological rationale for the use of operant conditioning and high-fidelity, and unimodal neurofeedback.

## 7. Limitations

While our findings provide a new model for MI, several limitations must be acknowledged:

### 7.1 Task Signal mismatch

The primary limitation is the mismatch between the instructed ‘sit-to-stand’ task and the decodable C3/C4 hand-area signal (Result 5.1). This raises unanswerable questions about task compliance. Our conclusions are therefore specific to the C3/C4-centric network that participants engaged, not necessarily the ‘sit-to-stand’ network itself.

### 7.2 Spatial resolution and Source-Space Analysis

Our network analysis was restricted to a 17-channel subset. This spatial downsampling limits our ability to make claims about whole-brain network organization. Our source-space analysis (Result 5.5) was a supplementary check, not a full validation, as reliable source reconstruction from 17 channels is extremely limited.

### 7.3 Putative Effective Connectivity (Granger Causality)

Our “Broadcasting M1” model is based on sensor-space GC and must be interpreted with extreme caution. GC is highly susceptible to confounds from volume conduction and common referencing, which can create spurious, non-physiological directional estimates (36, 37). Our analysis did not use methods robust to these confounds (e.g., the imaginary part of coherency) (35) and was not validated in source space. Furthermore, GC model order and data stationarity were not explicitly assessed (38). Therefore, the GC results (Table 4) are presented as an *exploratory, functional hypothesis*, not as a definitive measure of causal information flow.

### 7.4 Statistical Power

Our primary null finding (r = −0.054 with N=32) must be interpreted with respect to statistical power. An a priori power analysis (for α = 0.05, 80% power) indicates our sample of 32 participants was sufficient to detect a *large* correlation of r ≈ 0.48 or greater (39). Our 95% CI for the source-space correlation was wide ([−0.395, 0.300]). Therefore, while we can confidently rule out a large effect, our study was underpowered to detect a small or moderate relationship.

### 7.5. Lack of Null-Model Comparison

A significant methodological limitation of this study is that our graph metrics were not statistically validated against a null-model, such as a degree-preserving rewired network (21). Therefore, while we identified topological structures like “hubs” (e.g., F3/Fz) and “gateways” (e.g., FC1/P3), we cannot formally claim that these metrics are statistically significant or that the network has a non-random, “small-world” topology. This must be a priority for future validation.

### 7.6. Methodological and Population Limitations

Our results are also based on a single 10% density threshold, and graph metrics are known to be sensitive to this choice (30). A further limitation is that trial counts and rejection rates per subject were not available for this analysis. Moreover, the preprocessing pipeline did not include manual ICA component rejection, relying instead on the amplitude threshold, which may have allowed non-stereotypical artifacts to remain. Finally, our cohort consisted of healthy, young, BCI-naive individuals, and these findings may not generalize to clinical or expert-user populations.

## 8. Future Directions

This study opens several new avenues for investigation:

1. **Task Compliance:** Replicating this study with a simple left-vs-right hand task to confirm if the C3/C4-centric network and the null correlation persist.
2. **Longitudinal BCI Learning:** The most critical follow-up is a longitudinal study. How does the local-global relationship change *as* an individual learns to control a BCI? We hypothesize that learning *strengthens* this dissociation, as the user learns to isolate and “tame” the local C3 signal.
3. **Multimodal Validation:** Our “Broadcasting Motor Cortex” model is a strong, testable hypothesis. It should be validated using methods with superior spatial resolution, such as high-density EEG (64+ channel), fMRI with Dynamic Causal Modeling (DCM), or with perturbative methods, such as applying TMS (4) to F3 to see if it disrupts “monitoring” without disrupting the core M1 “signal.”
4. **Clinical Translation:** This entire analysis pipeline should be applied to clinical populations. Do stroke patients with M1 lesions fail at BCI because their “broadcaster” is broken? Do patients with PFC lesions (impaired “monitors”) fail because they cannot *sustain* the MI state? This framework provides a new, systems-level diagnostic tool for BCI rehabilitation.

## 9. Conclusion

We have demonstrated that the motor imagery network—in this case, a C3/C4-centric network robustly recruited for a postural task—is a sophisticated system with stable prefrontal hubs. We provide evidence that the local decoding accuracy of a BCI is functionally dissociated from the global efficiency of the brain network during task execution (r = −0.054, p = 0.769). This clarifies the neurophysiology of BCI skill, suggesting that while efficient baseline networks may predict BCI aptitude (16, 17), task-active global integration is not a correlate of performance. While our study was underpowered to detect small or moderate effects (see Section 7.4), this robust null finding for a large effect indicates that local BCI skill and *task-active* global integration are not strongly coupled.

## 10. Conflict of Interest Statement

The authors declare that the research was conducted in the absence of any commercial or financial relationships that could be construed as a potential conflict of interest.

## 11. Funding

The authors did not receive any financial support for the research, authorship, or publication of this article.

## 12. Acknowledgments

This study utilized the publicly available OpenNeuro dataset ds005342. The authors express their gratitude to the original data contributors (Nayid Triana-Guzman, Alvaro D Orjuela-Cañon, Andres L Jutinico, Omar Mendoza-Montoya, and Javier M Antelis) for their commitment to open science, which made this work possible (22). The authors acknowledge the use of Google’s Gemini AI model to assist in generating, refining, and debugging code for data preprocessing and analysis. The model was also used to assist with paraphrasing and improving language clarity in the manuscript. All outputs were reviewed and validated by the authors, who take full responsibility for the final content.

## 13. Data Availability Statement

The EEG dataset (ds005342) analyzed in this study is publicly available in the OpenNeuro repository, accessible at https://doi.org/10.18112/openneuro.ds005342.v1.0.3 (22). The analysis scripts generated in this study, along with an environment file for reproducibility, are publicly available on GitHub at https://github.com/draahayon-bd/MI-BCI-Network-Analysis. These scripts were developed with partial assistance from Google’s Gemini AI model.

## Notes

### Competing Interest Statement

The authors have declared no competing interest.

